# Metabolically flexible microorganisms rapidly establish glacial foreland ecosystems

**DOI:** 10.1101/2025.02.19.639150

**Authors:** Francesco Ricci, Sean K. Bay, Philipp A. Nauer, Wei Wen Wong, Gaofeng Ni, Luis Jimenez, Thanavit Jirapanjawat, Pok Man Leung, James A. Bradley, Vera M. Eate, Montgomery Hall, Astrid Stubbusch, Beatriz Fernández-Marin, Asunción de los Ríos, Perran L.M. Cook, Martin H. Schroth, Eleonora Chiri, Chris Greening

## Abstract

An overriding question in ecology is how new ecosystems form. This question can be tested by studying colonisation of environments with little to no pre-existing life. Here, we investigated the functional basis of microbial colonisation in the forelands of a maritime Antarctic and an alpine Swiss retreating glacier, by integrating quantitative ecology, genome-resolved metagenomics, and biogeochemical measurements. Habitat generalists and opportunists rapidly colonize both forelands and persist across soil depth and decadal chronosequences that serve as proxies for temporal community dynamics. These microbes are metabolically flexible chemotrophic aerobes that overcome oligotrophic conditions by using both organic and inorganic compounds, including atmospheric trace gases and sulfur substrates, for energy and carbon acquisition. They co-exist with metabolically flexible early-colonising opportunists and metabolically restricted later-colonising specialists, including photosynthetic Cyanobacteria, ammonia-oxidising archaea, and obligate predatory and symbiotic bacteria, that exhibit narrower habitat distributions. Analysis of 589 species-level metagenome-assembled genomes reveals early colonisation both by generalists and opportunists is strongly associated with metabolic flexibility. Field- and laboratory-based biogeochemical measurements reveal the activity of metabolically flexible microbes rapidly commenced in the forelands. Altogether, these findings suggest primary succession in glacial foreland soils is driven by self-sufficient metabolically flexible bacteria that mediate chemosynthetic primary production and likely provide a more hospitable soil environment for subsequent colonisation.

## Introduction

Environments with little or no preexisting life can serve as natural laboratories to understand how complex communities establish and thrive^1–5^. Pioneer microorganisms (i.e. primary colonizers) establish such systems, performing key roles such as primary production, nutrient recycling, weathering, and detoxification, thereby increasing habitability and supporting secondary colonization of diverse species^6–9^. Once microbial communities are established, cryptogams, vascular plants, and animals can sometimes colonize^9^. Over time, the assembly of microbial communities is governed by the interplay between stochastic (including dispersal from air and water flow) and deterministic (including selection due to biotic and abiotic factors) processes^10,11^. As pioneer ecosystems are typically carbon- and energy-limited, primary producers are particularly important ecosystem engineers^12^. Accordingly, most classical studies on pioneer microorganisms have focused on photosynthetic microbes such as Cyanobacteria^13–15^. Primary producers also include chemosynthetic microbes that use lithic substrates (e.g. NH ^+^, Fe^2+^, and S^2−^)^16,17^ and trace gases (e.g. H_2_, CO, and CH)^18,19^ to drive aerobic respiration and carbon fixation. While photosynthetic microbes are often metabolically constrained by light and water availability, many chemosynthetic microbes are highly flexible, for example alternating between inorganic and organic substrates in response to fluctuating environmental conditions and resource availability, and thus may be able to better tolerate the harsh and variable conditions of many pioneer ecosystems^20–22^. Our discovery of aerotrophy, i.e. the use of atmospheric trace gases to drive aerobic respiration and carbon fixation particularly in oligotrophic environments (e.g. Antarctic deserts)^48–50^, suggests that the atmosphere may be a particularly dependable energy supply for microbial pioneers that may enable continuous primary production during ecosystem establishment. To date, multiple studies have investigated compositional changes and ecological dynamics of microbial communities during colonization and succession^11,23^. However, the functional traits and biogeochemical processes that enable microbial colonizers to colonize remain unresolved.

Glacial forelands are ideal systems to study ecosystem formation^24^. The retreat of land-terminating glaciers, as accelerated by anthropogenic global warming, produces forelands that are gradually colonized. The primary succession of these forelands can be studied using a chronosequence approach (assuming space-for-time substitution^25^), where sites of increasing distance away from the retreating glacier front represent increasing soil ages^9,26–29^. Recently deglaciated lands are harsh environments for microbial colonizers, typically characterized by low organic carbon and nitrogen, high irradiance, and large temperature fluctuations, which impose strong selective pressures on colonization^9,27,30^. These pressures can be especially extreme in places that are already at the fringe of life, such as terrestrial Antarctica. In such systems, early colonization processes involve a trade-off between surviving these physicochemical pressures and exploiting available resources^30^. Initial colonizers may disperse from various sources: for instance, ice-dwelling microbes can persist on deglaciated soil^31^, while others can be airborne or transported through hydrologic flow^32,33^. Regardless of their origin, selected microbes rapidly colonize and mediate biogeochemical cycling in newly exposed soil, facilitating secondary colonization^9,34^. Previous studies have begun to use metagenomics to gain insight into the functional strategies supporting microbial colonization^27,38,39^. For example, they have suggested that primary colonizers overcome nitrogen limitation in forelands through a multiplicity of strategies, including assimilation of inorganic nitrogen, degradation of organonitrogen compounds, and nitrogen fixation^27,30,40^. While certain biogeochemical processes are conserved throughout foreland chronosequences, later successional stages may result in the emergence of other metabolisms, such as methane oxidation and nitrification^19,27,40,41^. Yet despite this emerging knowledge base, we lack a consolidated understanding of the relationship between the ecological dynamics, functional traits, and biogeochemical activities driving microbial colonization and subsequent succession, including the metabolic breadth of pioneer microbes and roles of different primary production strategies^3,24,29^.

Ecological theory, originally developed for plant communities, provides a quantitative framework to examine colonization and succession dynamics^42,43^. A central theory is that habitat generalists (i.e. organisms that can withstand a wide range of environmental conditions and resource availabilities) are early colonizers, while specialist taxa (i.e. organisms exhibiting narrower habitat preferences and with stricter resource requirements) generally increase during succession^44,45^. This concept, together with trait-based frameworks^44^, offers a baseline to examine microbial primary succession from a functional standpoint. However, it is important to note that microbes generally exhibit greater metabolic flexibility than plants and animals, with many able to use multiple carbon and energy sources either simultaneously (mixotrophy) or alternatively^21,46^. Most can also meet energy needs under resource limitation through continuous energy harvesting using trace gases or sunlight (*via* rhodopsins and photosystems) while in dormant or slow-growing states^47^. Therefore, it is possible that microbial communities do not conform to the classic macroecological succession dynamics because of their high degree of flexibility, which could provide an ecological advantage from early to late successional stages^47^. Given these considerations, we hypothesized that the dominant pioneers in glacial forelands would be metabolically flexible bacteria that overcome nutrient limitation by conserving energy and fixing carbon using atmospheric trace gases. Here we provide a system-wide understanding of the community and functional dynamics underpinning colonisation of two glacial forelands, by integrating gene- and genome-centric metagenomics with biogeochemical measurements and ecological theory. Our study focused on two contrasting sites. Hurd Glacier on Livingston Island, Antarctica is a remote maritime glacier, terminating over the Southern Ocean, that forms part of the 10 km^2^ Hurd Peninsula ice cap alongside Johnsons Glacier^51^. Griessfirn Glacier in Switzerland is also a small (∼5 km^2^) glacier, but is an alpine glacier at 2400-3300 meters above sea level^52^, and is more likely to be subject to external resources input and organism colonization / invasion due to its central position in the Swiss Alps. Our findings provide a comprehensive understanding of microbial metabolic strategies that enable the colonization and establishment of ecosystems from their earliest stages of succession to communities that have persisted for over a century.

## Results and Discussion

### Microbes rapidly and deterministically colonize foreland chronosequences

We sampled surface foreland soils of a maritime Antarctic and an alpine Swiss glacier, covering chronosequences of 21 and 127 years since deglaciation, respectively **(Fig. 1a)**, as well as depth profiles (from 0 to 50 cm) for the Swiss soils. Geochemical analyses showed that the foreland soils have low organic carbon (av. 0.14 ± 0.02% Antarctic, 0.57 ± 0.24% Swiss) and nitrogen (typically <0.02%), with low salinity and moderate pH **(Table S1)**. Thus as anticipated^9^, these newly deglaciated soils are highly oligotrophic especially at the Antarctic site and therefore are challenging substrates for microbial colonisation. Yet complex microbial communities inhabited even the most recently deglaciated soils **(Fig. 1b)**. Our study design does not allow tracking of the origins of the microbes present, but potential sources include snow and airborne dispersal, glacier meltwater transport and deposition, and microbes that previously inhabited the subglacial environment that became exposed by ice retreat^31,53,54^. Microbial abundance (16S rRNA gene copy number) increased on average tenfold across the chronosequences at both glaciers **(Fig. 1c)**. Similarly, microbial richness increased along the chronosequences, with Antarctic soil showing the sharpest increase from recently deglaciated (av. 196 amplicon sequence variants, ASVs; Shannon index 4.33) to more mature soils (692 ASVs, Shannon index 5.71) **(Table S2; Fig. 1d)**. Despite this, microbial composition of the two sites (based on β-diversity ordinations) were distinct, with strong differentiation by site location (*p* < 0.001) and to a lesser extent soil age (*p* < 0.001) **(Table S2; Fig. 1e)**. Following colonisation, we observed an increase in community similarity in later stages of succession **(Fig. 1e)**, as previously observed in Antarctic forelands^28^. The communities were also strongly structured by depth, decreasing in abundance (av. 128-fold) and richness (av. 4.9-fold) from surface (0-10 cm) to deep (40-50 cm) samples **(Fig. 1a & 1b; Table S2)**. These patterns, consistent with previous studies^28,55,56^, suggest that foreland microbial communities may respond to common successional dynamics^57^.

**Figure 1.**
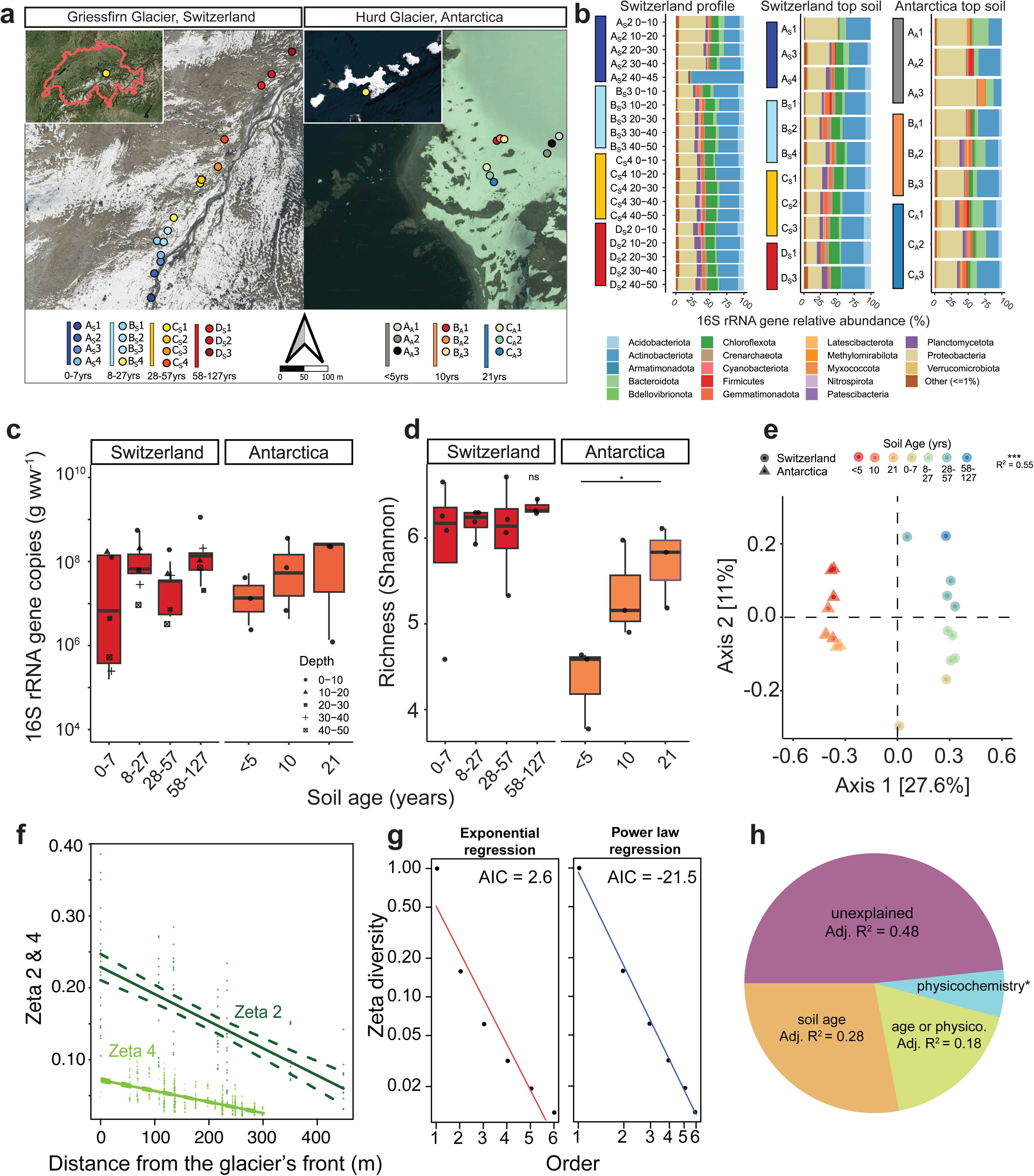
Deterministic processes drive microbial community assembly in glacial forelands. **(a)** Satellite imagery showing the geographic setting of the glacier forelands (Switzerland left, Antarctica right) studied at local and regional level. Sampling design, major environment features, study, scale and sites are shown. **(b)** Stacked barchart showing phylum-level community structure at the sample level. Vertical-coloured bars show the sample clusters according to soil age. **(c)** Boxplot showing 16S rRNA gene copy number per gram wet weight (g_ww_) for soils at multiple depths and according to location and soil age. **(d)** Boxplot showing microbial community richness (Shannon diversity index) for topsoil (0-10 cm), according to location and soil age. Statistical differences across groups were assessed using ANOVA, with a significance threshold of *p* = 0.05. **(e)** Unconstrained Principal Coordinate Analysis (PCoA) showing beta diversity (Bray-Curtis dissimilatory index) of topsoil (0-10 cm) and compositional differences in community structure according to location and soil age. (**f**) Zeta distance decay relationship showing community turnover across the Swiss chronosequence. The number of ASVs shared between two sites (ζ_2_) and four sites (ζ_4_) with distance are compared. (**g**) Zeta decline of the Swiss chronosequence modelled by exponential and power law regressions, associated with stochastic and deterministic processes respectively, with Akaike Information Criterion (AIC) scores. (**h**) Variation partitioning of zeta diversity across two sites (Zeta orders) of the Swiss chronosequence, explaining variation due to distance and physicochemical factors (*adj. R² = 0.059).

Communities at both glacial forelands were dominated by common soil phyla, especially Proteobacteria and Actinobacteriota **(Fig. 1b)**, in line with previous studies of receding glaciers^27,58–60^. This was also reflected at finer resolution, with Proteobacteria (e.g. *Devosia*, *Methylotenera*) and Actinobacteriota (e.g. *Pseudarthrobacter*, *Gaiella*) accounting for two thirds of the hundred most abundant ASVs and half of the hundred most widespread ASVs across the two sites **(Table S2)**. The Swiss samples also contained two highly abundant archaeal ASVs (av. 1.6% community) that affiliated with the family Nitrososphaeraceae, known to harbour cold adapted ammonia oxidizers^61^. Diverse photosynthetic Cyanobacteria were also detected, spanning 59 ASVs, though they were at low relative abundance (av. 0.13%) and absent from the entirely deglaciated Antarctic sites **(Table S2)**; an exception was one Swiss topsoil (25-50 years old) where a single ASV from the filamentous genus *Microcoleus* comprised 2.9% of the community, suggesting these habitat specialists can thrive in certain established soils **(Fig. 1b)**. Concordantly, chlorophyll *a* from Cyanobacteria and microalgae was minimal across the forelands of both glaciers (av. 0.52 µg g_soil_^-1^ Swiss; av. 1.30 µg g_soil_^-1^ Antarctic) but increased progressively with soil age **(Table S2)**. Similarly, some methanotrophs (e.g. *Methylomonas* and *Methyloglobulus*) were present but at low occupancy. The Antarctic sites also contained high but variable levels of the obligate predator Bdellovibrionota (av. 1.8%) and obligate symbiont Patescibacteria (av. 2.3%), in support of our previous observations in Antarctic soils^50^. To substantiate these inferences, we calculated specialization indices for each genus based on the coefficient of variance of their relative abundance across samples as previously described^21,62^ **(Table S2)**. This confirmed the co-existence of habitat generalists with broad occupancies (e.g. *Pseudarthrobacter* and *Gaiella*) alongside habitat specialists with narrow distributions (e.g. *Microcoleus* and *Methyloglobulus*) and those with intermediate specializations (e.g. *Methylotenera* and *Polaromonas*; present in all Antarctic samples but at highly variable abundance).

Zeta diversity was used to understand the patterns and drivers of community turnover along the Swiss glacial forelands. This new incidence-based metric compares the average number of taxa shared between multiple sites as the number of sites (zeta decline) or distance (zeta decay) increases^63,64^. Zeta diversity declined at a moderate rate, with an average of 23% of ASVs shared between two sites (ζ_2_) and 6.6% between four sites (ζ_4_) **(Fig. S1)**, and the coefficient of distance decay was threefold higher for two-way compared to four-way comparisons **(Fig. 1f)**. Together, these zeta diversity and habitat specialization calculations suggest that most microbes inhabiting these forelands are habitat specialists that undergo rapid turnover, but co-exist with many habitat generalists that persist across the transect and are disproportionately abundant **(Fig. S1)**. Zeta decline best fit a power law regression **(Fig. 1g)**, suggesting deterministic not stochastic factors primarily drove the composition of these sites at time of sampling. Variation partitioning analysis suggested community turnover was strongly driven by soil age (28%), soil physicochemistry (5.9%), and the interaction of these factors (18%) **(Fig. 1h).** Overall, these findings suggest both glacier forelands are rapidly and successively colonized by microorganisms through deterministic processes, potentially linked to factors such as nutrient loading, microbial interactions, and soil stabilisation as the glacier retreats.

### Metabolically flexible microorganisms drive glacial foreland colonisation

We used genome-resolved metagenomics to differentiate the metabolic traits of habitat generalists and specialists in the community. For the Antarctic and Swiss glacier forelands respectively, 367 and 222 species-level (ANI: 0.95) metagenome-assembled genomes (MAGs) were assembled, spanning 19 and 17 phyla and capturing 33% and 10% of reads, respectively **(Fig. 2a)**. The specialization index for each MAG was calculated based on read mapping to the topsoil metagenomes, with MAGs classified as habitat generalists or specialists if their specialization indices were in the lower or upper quartiles respectively **(Table S3)**. These MAG-based classifications were generally concordant with the amplicon-based classifications, with clear differentiation of habitat generalists (e.g. Actinobacteriota such as *Gaiella*, Proteobacteria such as *Ramlibacter* among numerous novel genera) from habitat specialists (e.g. photosynthetic *Microcoleus*, methanotrophic *Methyloglobulus*, Bdellovibrionota and Patescibacteria). However, it also revealed that genus-level classifications sometimes obscured patterns by lumping species with different specialization indices, for example in the case of *Polaromonas* with six MAGs spanning the full spectrum of habitat generalists to specialists **(Table S3)**. The metabolic versatility of each MAG was inferred based on whether they encoded 56 signature metabolic genes, including the primary dehydrogenases for organotrophy and lithotrophy, the terminal reductases for aerobic and anaerobic respiration, and the signature enzymes for carbon fixation, photosystem- and rhodopsin-based phototrophy, and nitrogen fixation. The habitat generalist MAGs exhibited higher metabolic flexibility (encoding on average 8.8 and 7.5 signature metabolic genes in the Antarctic and Swiss datasets respectively) compared to the habitat specialist MAGs (6.5 and 6.2 respectively). This provides the first quantitative support for the hypothesis that metabolic flexibility enhances niche breadth and in turn promotes habitat generalism of microorganisms.

**Figure 2.**
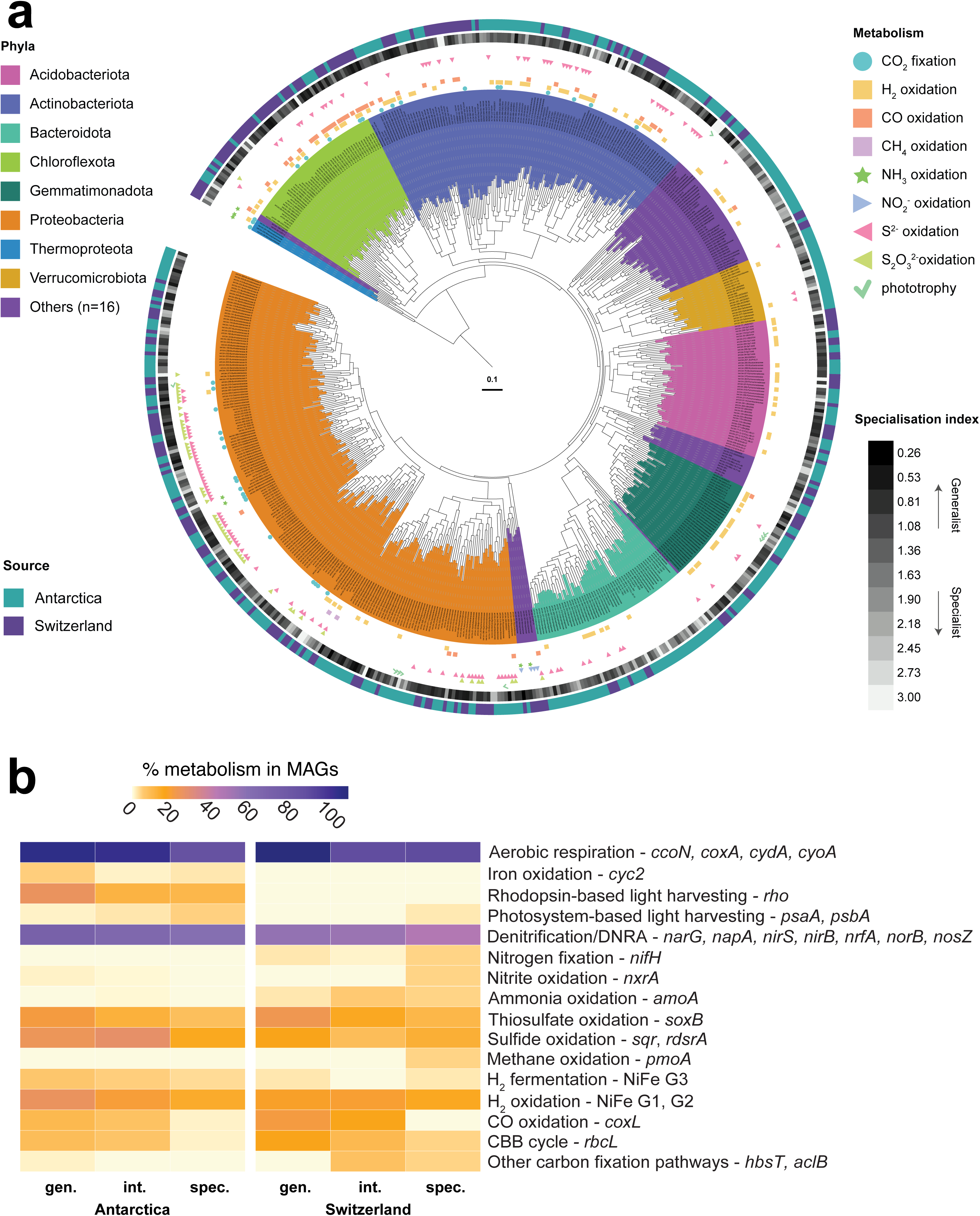
Metabolic traits differentiate habitat generalists and specialists in glacial forelands. **(a)** Phylogenomic tree showing the evolutionary history, metabolic capabilities, and habitat specialization of the 589 glacial foreland metagenome-assembled genomes (MAGs) spanning 24 phyla. The archaeal and bacterial tree was constructed using the maximum likelihood method, with LG+F+G4 substitution model with 1,000 iterations of ultrafast bootstrapping and midpoint rooting. Clades are coloured according to phylum-level taxonomy and symbols indicate the potential of each MAG to mediate key energy and carbon acquisition processes. The heatmap on the inner concentric ring represents the specialization index, while the outer concentric ring’s colours denote the source of each MAG (Swiss vs Antarctic). **(b)** Metabolic capabilities of MAGs identified as habitat generalists (gen.), intermediates (int.), or specialists (spec.) at the Antarctic and Swiss glaciers.

As expected from the geochemical analyses of these oxygenated carbon-poor soils **(Table S1)**, the MAG-based metabolic annotations suggest both forelands are limited primarily by energy rather than by oxidant and nitrogen supply (**Table S3)**. Almost all bacterial MAGs are predicted to mediate respiration using organic compounds as electron donors and oxygen as an electron acceptor, with much of the community also capable of one or more denitrification steps (65% MAGs; adjusted for MAG completion; **Table S3-4**). There was much capacity for microbes to also input electrons from inorganic and one-carbon compounds from both lithic sources, namely sulfide (33% MAGs), thiosulfate (16%), ammonia (1.9%), nitrite (0.9%), iron (1.7%), and potentially formate (42%), as well as the atmospheric trace gases hydrogen (22%), carbon monoxide (9.8%), and methane (0.43%; **Table S4)**. 10% of MAGs spanning five phyla were predicted to mediate carbon fixation, predominantly using chemosynthetic lineages of RuBisCO, suggesting many are self-sufficient primary producers that use electrons from atmospheric gases and lithic substrates increasing the organic carbon pool of the system. For instance, at the Antarctic foreland, an abundant proteobacterial MAG from the genus *Polaromonas* co-encoded CO dehydrogenase, thiosulfohydrolase, and reverse dissimilatory sulfite reductase with a form ID RuBisCO, indicating the capacity to use both atmospheric and lithic electron donors. Despite the minimal capacity for oxygenic photosynthesis, 12% of MAGs spanning seven phyla were predicted to be capable of photoheterotrophic growth by harvesting light using rhodopsins or photosystem II. Only four MAGs encoded nitrogenases, suggesting these soils were more strongly carbon-than nitrogen-limiting. Though the metabolic capabilities of the MAGs reconstructed from the two glaciers were similar, an exception was the capacity for phototrophy, which was encoded by 54 Antarctic MAGs (all predicted photoheterotrophs) compared to one Swiss MAG (the cyanobacterium *Microcoleus*). Conversely, the capacity for nitrification was higher in the Swiss MAGs. Overall, these data suggest that the primary colonizers of glacier forelands exhibit significant metabolic flexibility, with aerotrophy and classical lithotrophy emerging as dominant processes, likely enabling their adaptation to these oligotrophic settings and providing resources for the establishment of secondary colonizers.

The habitat generalists and specialists differed in their energy and carbon acquisition strategies across the two glaciers **(Fig. 2b)**. For example, RuBisCO genes for carbon fixation through the Calvin-Benson-Bassham cycle were 5.6-fold more abundant in habitat generalists and were almost exclusively associated with chemosynthetic bacteria **(Table S3)**. Of these RuBisCO-encoding generalists, 54%, 37%, and 29% were capable of oxidising sulfur compounds (*via* sulfide-quinone oxidoreductase, reverse dissimilatory sulfite reductases, and thiohydrolase; e.g. *Ramlibacter*), atmospheric H_2_ (*via* group 1h and 1l [NiFe]-hydrogenases; e.g. Chloroflexota CSP1-4), and atmospheric CO (*via* form I CO dehydrogenases; e.g. *Pseudonocardia*). The generalist MAGs also had a greatly enhanced capacity for oxidation of inorganic compounds compared to specialists, most strikingly CO (25-fold), but also H_2_ (1.7-fold), sulfide (1.7-fold), thiosulfate (2.7-fold), and iron (2.7-fold), as well as rhodopsin-based light harvesting (2.9-fold). This is consistent with the hypothesis that habitat generalists are metabolically flexible, often self-sufficient microorganisms with the capacity to continuously harvest energy and mediate chemosynthesis. Conversely, the genes for photosynthesis, methanotrophy, and nitrification were encoded primarily by metabolically restricted habitat specialists. For example, two seemingly obligately methanotrophic MAGs were retrieved from the Swiss forelands, classified as *Methyloglobulus* and *Methyloligotrophales,* the latter a key aerotrophic clade in desert and cave ecosystems^48^. A notable exception was a Nitrosopiraceae MAG (genus Palsa-1315, also known as Clade B Comammox *Nitrospira*), predicted to be capable of complete nitrification, that was ubiquitous across the Swiss transect and among the 15 most abundant MAGs, and hence can be considered a metabolically restricted pioneer primary producer. Collectively, these findings lend further support to the theory that habitat generalists are usually more metabolically versatile and hence can use a broader range of resources than habitat specialists^21^.

### Ecologically and functionally distinct microbes vary in abundance over soil age

We next sought to understand how the ecological traits and metabolic capabilities of the microbial communities varied across the chronosequence. First, we mapped the relative distributions of habitat generalists and specialists with soil age based on both the amplicon and metagenome data. While the absolute abundance of both generalists and specialists increased during succession, their relative proportions shifted. Based on the amplicon datasets, while habitat generalists were abundant across all samples, they formed a lower proportion of the community in the initially exposed soils (av. 46% community) and became dominant in all other soils (av. 66% community) in both forelands **(Fig. 3a).** This is in line with the observed stabilisation of community structure with time **(Fig. 1e)**. Similarly, for the metagenome data, habitat specialists were most enriched during the initial colonisation **(Fig. 3b)**. Such observations were contrary to our initial hypothesis, as we expected habitat generalists to drive initial colonisation and habitat specialists to be most dominant in the oldest soils. While a slight increase in the relative abundance of habitat specialists was observed in the oldest soils compared to the medium age soils, their proportion was highest in the youngest soils overall **(Fig. 3a & 3b)**. Nevertheless, these observations remain concordant with broader macroecological theory: in line with our recent modification of Grime’s theory^43^ (CSO: competitor / stress tolerator / opportunist framework), the early-colonizing specialists are likely opportunists primed to rapidly colonize new environments by exploiting available resources released during deglaciation, but become outcompeted by stress-tolerant generalists and later-colonising specialists (including competitors) at later successional stages.

**Figure 3.**
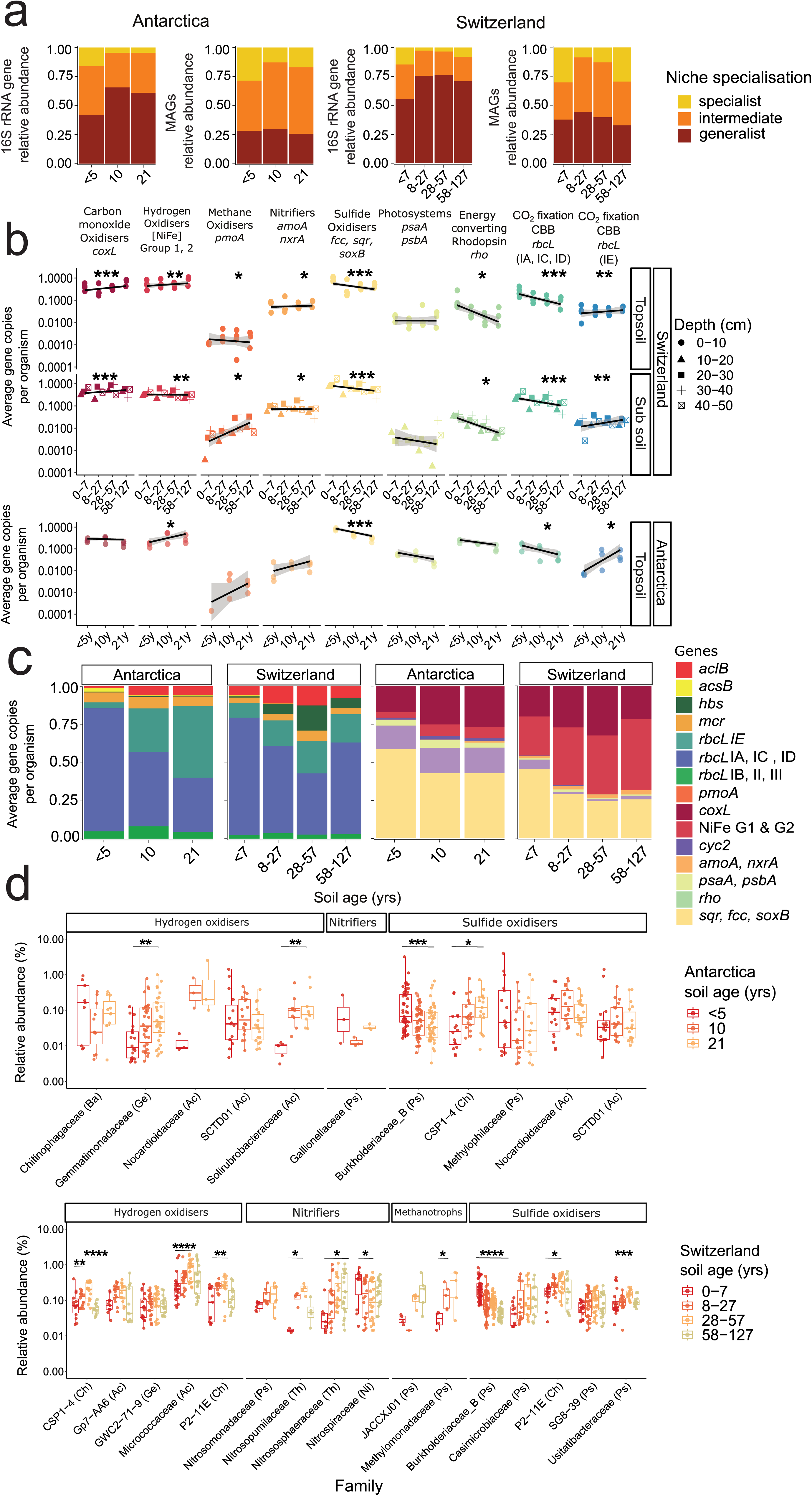
Microbial habitat classes and metabolic capabilities vary across the glacial forelands. **(a)** Proportion of specialist, intermediate and generalist microbes classified based on 16S rRNA gene amplicon sequencing and metagenome-assembled-genomes (MAGs) across the Antarctic and Swiss glacier forelands. **(b)** Dot plots showing average gene copy per organism of topsoil (0-10 cm) based on short read analysis of key metabolic marker genes, with a fitted Generalized Linear Model (GLM) for the Swiss and Antarctic glaciers. The fitted line represents the linear relationship under a Gaussian error distribution, with shaded ribbons showing 95% confidence intervals. Asterisks represent the lowest *p* value among coefficients of soil ages (GLM), derived from Wald test-based inference for each coefficient. The same is shown for subsoil samples (10-50 cm) of the Swiss glacier. **(c)** Proportion of carbon fixation and energy conversion marker genes across the Antarctic and Swiss glacier forelands based on short read analysis. **(d)** Boxplots showing the relative abundance of selected MAGs predicted to be capable of oxidising lithic and atmospheric substrates across the Antarctic and Swiss glacier forelands. Statistical differences among age groups within MAG families were assessed using the Kruskal-Wallis test. A Dunn’s post-hoc test with Bonferroni correction was applied to identify significant pairs, with asterisks denoting significance.

These predictions are supported by analysis of variations of specific taxa with age. Proteobacterial habitat specialists dominate the newly exposed soils, for example with a *Methylotenera* MAG and a SURF-13 MAG comprising 1.9% and 1.5% of the community (based on metagenomic read mapping) respectively in the earliest Antarctic soils (0 – 5 yrs), but comprise less than 0.1% of the community in more established soils (21 yrs) **(Fig. 3c; Table S4)**. Such taxa are also quantitatively more abundant in early rather than late soils (based on 16S rRNA gene copy number normalisation). A similar pattern was observed for the Swiss soils, with MAGs from multiple Burkholderiaceae genera (e.g. *Methylibium*, *Polaromonas*, CADEEN01) abundant in the earliest soils (0 – 7 yrs) but either minimal or undetected in the oldest soils (58 – 127 yrs) **(Fig. 3c; Table S4)**. Importantly, these opportunistic primary colonizers are still metabolically flexible, capable of using lithic substrates (e.g. sulfide) and often atmospheric hydrogen to support lithoheterotrophic and sometimes lithoautotrophic growth. The later-colonizing habitat specialists, for example ammonia oxidizers (e.g. Nitrosomonadaceae, Nitrososphaeraceae) and the aforementioned methanotrophs, show the opposite trend and fill metabolic niches unexploited by other colonizers. In contrast, the dominant habitat generalists were also present in the earliest soils, but typically increased modestly in relative abundance and greatly in absolute abundance across each chronosequence; these trends are particularly pronounced for the most abundant lineages of Actinobacteriota (e.g. Nocardioidaceae, Micrococcaceae) and Chloroflexota (e.g. CSP1-4, P2-11E; **Fig. 3c)**, each of which is predicted to be metabolically flexible.

This shift in generalist *versus* specialist distributions during foreland succession is also reflected by differences in energy and carbon acquisition genes. Based on community-wide profiling using metagenomic short reads (as opposed to MAG-based analysis; **Fig. 2**), most microbial cells inhabiting the forelands can use inorganic electron donors, including sulfide (av. 32% Swiss topsoil cells, 48% Antarctic cells), H_2_ (51%, 36%), CO (35%, 29%), and/or ammonium (3.2%, 0.65%) with considerable capacity also for rhodopsin-based light harvesting (4.1%, 21%) and carbon fixation (21%, 19%) **(Table S3; Fig. 3b)**. In line with being encoded primarily by opportunistic early colonizers, the proportion of the community capable of sulfide oxidation strongly and consistently decreased with soil age for both forelands (*p* < 0.001), whereas hydrogenases and CO dehydrogenases modestly increased with soil age (*p* < 0.01), in agreement with their association primarily with stress-tolerant habitat generalists **(Fig. 3b)**. Rhodopsin and photosystem genes also decreased with soil age, suggesting that light harvesting can be adaptive for initial colonisation, though this is a less common strategy than lithotrophy. Carbon fixation pathways shifted with soil age in both forelands. The RuBisCO lineages typical of classical chemolithoautotrophs (i.e. form IA, ID and to a lesser extent IC)^65,66^ dominate in both forelands in the earliest soil ages **(Fig. 3c & 3d)** and, based on genome-resolved analysis, are primarily encoded by sulfide-oxidizing Proteobacteria and metabolically flexible Chloroflexota **(Table S3)**. In contrast, the form IE RuBisCO typical of aerotrophs^48,49,67^ are encoded exclusively by diverse Actinobacteriota MAGs and become dominant in more established soils **(Fig. 3c & 3d; Table S3)**. Random forest analysis emphasised that soil age since deglaciation is a key driver of the distribution of these genes, especially for sulfide oxidation, in line with a key role of this process in early colonization of carbon-poor soils, alongside other factors such as pH and nutrient concentrations **(Fig. S2)**. Using the Swiss samples, we also examined variations in the levels of metabolic genes with soil depth; while lithic and gaseous energy sources were predicted to be important throughout the soil profiles **(Fig. 3b)**, as expected, phototrophy genes decreased and anaerobic metabolism genes (e.g. for acetogenesis, dehalorespiration, hydrogenogenic fermentation) increased substantially with soil depth **(Fig. 3b; Table S3)**.

### Foreland microbial communities oxidize atmospheric and lithic substrates during colonisation

Finally, we conducted biogeochemical analyses to both validate metagenomic predictions and determine energy dynamics of foreland ecosystems during establishment. Based on physicochemical analysis of the soils **(Fig. 4a)**, ammonium, sulfur compounds, and organic carbon are present in sufficient amounts to support lithotrophic and organotrophic growth of high-affinity pioneer microbes; levels of these compounds were low, especially in the Antarctic forelands, except for the sulfur-enriched soils closest to the Swiss alpine glacier, with previous studies suggesting sulfur compounds accumulate due to glacial weathering and meltwater delivery^27,68,69^. Consistent with these observations, both Swiss and Antarctic soils oxidized sulfide in aerobic microcosm assays **(Fig. 4b)**; rates were highest at soils close to the glacier fronts, in line with their enrichment with sulfide-oxidizing Proteobacteria **(Fig. 3b)** and elevated substrate availability at the Swiss but not Antarctic site **(Fig. 4a)**. Reflecting the presence of high-affinity nitrifiers in the glacial forelands, microcosm experiments confirmed that ammonium was oxidized to nitrite and nitrate by the foreland communities under aerobic conditions **(Fig. 4c)**. Ammonium oxidation to nitrate occurred at similar rates throughout the Swiss foreland, in line with the presence of Nitrosopiraceae habitat generalists capable of complete nitrification. By contrast, nitrite accumulated in the microcosms with the early-stage Antarctic soils but was largely converted into nitrate in the later-stage soils **(Fig. 4c)**, consistent with the metagenomic data that nitrite-oxidizing and complete nitrifying bacteria are associated with later stages of succession **(Fig. S3)**. Together, these findings suggest lithic energy sources are used by foreland microbes to support aerobic respiration and likely primary production, with sulfide oxidation potentially serving as a particularly important driver of initial colonization.

**Figure 4.**
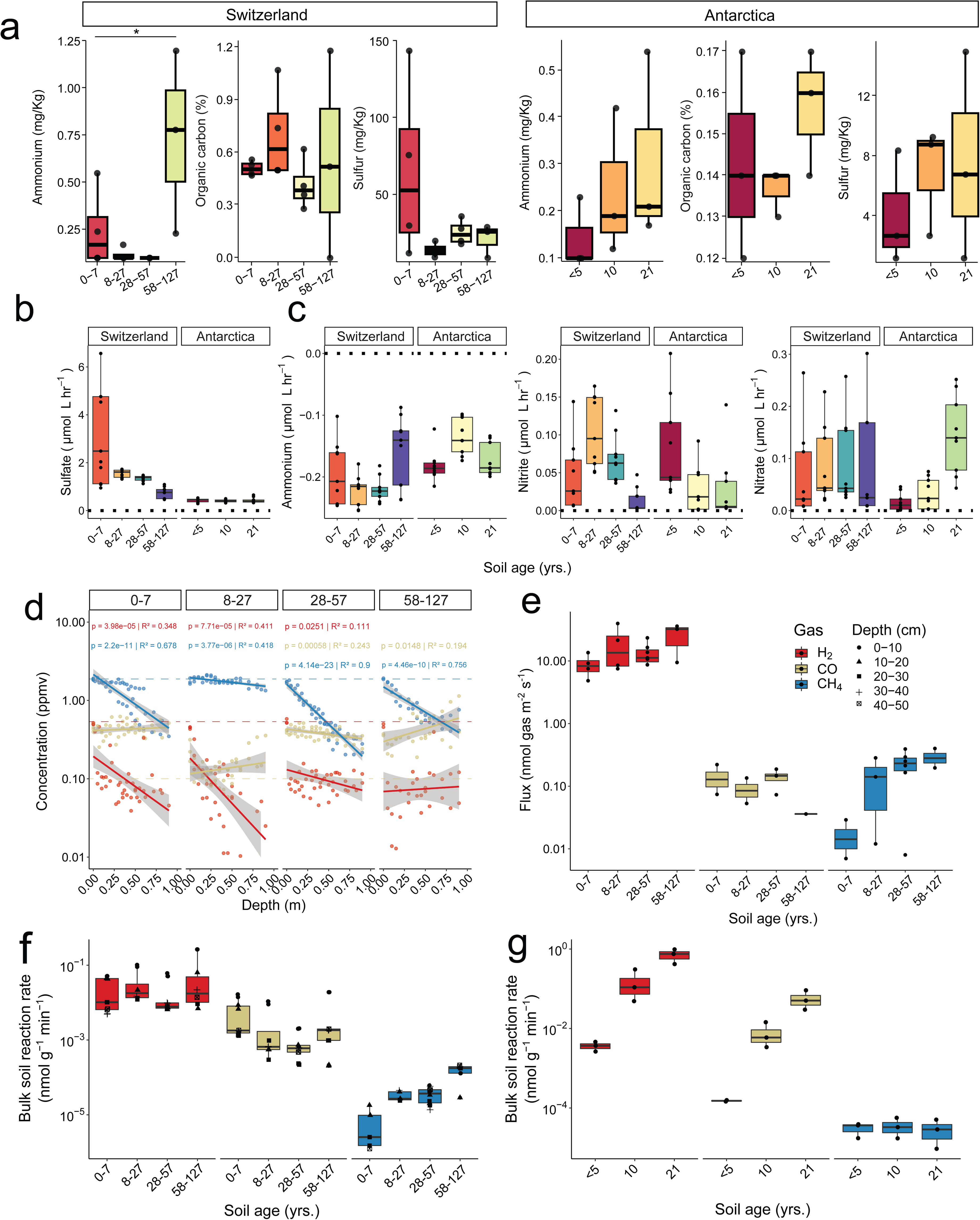
Trace gases and lithic compounds are major substrates that are differentially consumed across the chronosequences. **(a)** Boxplots show nutrient concentrations across the Swiss and Antarctic glacier forelands, with measurements taken from three biological replicates per age group. Statistical differences among age groups for each nutrient were determined using ANOVA or the Kruskal-Wallis test. **(b-c)** *Ex situ* oxidation rates of sulfur (b) and nitrogen (c) species were measured in microcosm experiments containing 10 g of soil and 200 mL of ultrapure water, supplemented with 50 µM of ammonium chloride or sodium sulfide. Positive values indicate accumulation, while negative values indicate uptake. Measurements were performed with at least three biological replicates per age group. **(d)** Dot plots illustrate *in situ* soil gas concentrations at Swiss sites relative to soil depth spanning ambient atmospheric conditions to deep subsoils (0-1 m). Linear models depicting significant relationships annotated with *p* values and R^2^ coefficients, with shaded ribbons showing 95% confidence intervals. **(e)** *In situ* soil–atmosphere gas fluxes of Swiss samples for each gas, with a minimum of two biological replicates per age group. **(f)** Bulk soil oxidation rates for each gas by soil age in Swiss samples, with different shapes indicating respective soil depths. Measurements were conducted on a minimum of five biological replicates per age group. **(g)** Bulk soil oxidation rates for each gas by soil age in Antarctic samples, measured with three biological replicates per age group.

To test our hypothesis that atmospheric trace gases are critical for ecosystem establishment, we also profiled the *in situ* levels and fluxes of trace gases in the Swiss soils (logistical constraints prevented equivalent analysis in Antarctic soils). H_2_, CO, and CH_4_ were present at the soil-atmosphere interface at mixing ratios typical of atmospheric averages (0.50 ± 0.04, 0.41 ± 0.12, 1.88 ± 0.14 ppmv respectively) and at variable levels at 1 m depth (0.074 ± 0.040, 0.50 ± 0.32, 0.56 ± 0.43 respectively); multiple factors influence the *in situ* levels of these gases, including gas exchange and diffusion rates in soil, biological consumption, and abiotic and biotic production processes, though the sharp depth-related decreases for H_2_ and CH_4_ suggest consumption by hydrogenotrophs and methanotrophs **(Fig. 4d)**. Consistently, *in situ* flux profiling revealed both gases were consumed, albeit at differential rates across the foreland. H_2_ was consumed at rapid rates across the whole foreland chronosequence with an average threefold increase from early to late deglaciated soils **(Fig. 4e),** in line with the gene- and genome-centric data. In contrast, uptake of CH_4_ established at later successional stages, increasing on average 17-fold from early to late deglaciated soil (Fig. 4a), supporting previous measurements^70^ and the observed distribution of methanotrophs and their marker genes **(Fig. 1 – 3)**. As with our previous investigations of organic soils^19^, we did not observe clear trends in CO levels or fluxes despite the widespread distribution of CO dehydrogenase and high *ex situ* CO oxidation activities **(Fig. 4f & 4g)**, with abiotic production within the flux chamber potentially obscuring trends. These findings support that trace gases are critical energy sources in these oligotrophic environments, with H_2_ oxidation emerging as an early-establishing, generalist-associated trait and CH_4_ oxidation as a later-establishing, specialist-associated one.

To help resolve to what extent atmospheric trace gases support foreland development, we measured their oxidation in Swiss and Antarctic samples using *ex situ* microcosms. Across all the samples, H_2_ was most rapidly consumed followed by CO and finally CH_4_ **(Fig. 4f & 4g)**, and oxidation rates generally declined with soil depth **(Fig. S3)**. For the Swiss samples, whereas CH_4_ oxidation greatly increased with soil age, H_2_ and CO oxidation occurred at consistent rates across the chronosequence **(Fig. 4f)**. In contrast, oxidation rates increased with soil age for the Antarctic soil, in part driven by an increased absolute abundance of H_2_ and CO oxidizers during succession. Although previous studies have shown forelands can be a sink for methane^71^, this is the first evidence of H_2_ and CO uptake in these communities. Building on these results, we investigated whether microbial communities could rely on H_2_, CO and CH_4_ for their energy demands **(Fig. S3)**. To do so, we used thermodynamic modelling to calculate the amount of power (J s^-1^ (W)) per cell that could be generated based on bulk trace gas oxidation rates **(Table S5)** and the total number of putative trace gas oxidizers detected per gram of dry soil **(Table S2 & S3)**. While rates varied between gases and sites, the average power per cell (3.76 × 10^-14^ W cell^-1^; range 3.33 × 10^-19^ to 2.22 × 10^-12^ W cell^-1^) is within the range to support the maintenance of all cells in the community (noting empirical measurements of maintenance energy range from 10^−12^ to 10^−17^ W cell^-1^ based on previous studies, with levels much lower in oligotrophs^72–75^). For some cells, the power derived from trace gas oxidation is sufficient to support mixotrophic or aerotrophic growth^76^. Altogether, these findings suggest that trace gases have a major role both in supporting colonization and subsequent development of a new terrestrial ecosystem.

## Conclusions

Despite their vastly different geographical locations and physicochemical properties, the primary colonizers of the Antarctic maritime and Swiss alpine glacier foreland were remarkably similar in functional capabilities. Rather than photosynthetic microbes, the most abundant and active ecosystem engineers appear to be versatile chemosynthetic bacteria that use atmospheric and lithic substrates. These findings highlight the central role of metabolic flexibility in shaping colonization and succession of new ecosystems. On one hand, metabolic flexibility provides opportunistic specialists with the capacity to rapidly exploit the varied resources liberated as glaciers retreat. On the other hand, this flexibility also provides habitat generalists with the capacity to adapt to the extensive physical, chemical, and biological changes that occur during different stages of ecosystem development. In both glaciers, there has been extensive selection for microbes that simultaneously or alternatively use organic, lithic, and atmospheric electron donors. Similar dynamics appear to influence the quantitative ecology, metabolic capabilities, and biogeochemical activities of the microbes inhabiting both forelands. Key differences include a more pronounced increase in microbial abundance and diversity over the Antarctic chronosequence, as well as potentially a stronger role for phototrophy in the Antarctic soils and ammonia- and sulfide-based lithotrophy in the Swiss soils. This convergence of findings suggests that microbial community assembly in glacier forelands is not primarily stochastic, but rather deterministic, as also supported by ecological and process modelling. The dynamic interplay between physicochemical conditions and biological interactions along the glacier chronosequences causes environmental filtering of communities, shifting specialist-generalist dynamics to enable co-existence of metabolically versatile habitat generalists with more restricted habitat specialists. Consistently, we observed increasing compositional stability over time, as also observed for other established glacier foreland microbial communities^55,77–79^.

In addition to deepening knowledge of the ecological dynamics of primary succession, by applying macroecological theory to microbial systems, this study also uncovers that atmospheric trace gas oxidation is a critical yet previously overlooked means for microorganisms to establish new ecosystems. This was confirmed through extensive metagenomic, flux, and activity measurements, as well as theoretical modelling. We also provide the first quantitative evidence that microbial metabolic flexibility is correlated with habitat generalism. Future work is needed to disentangle the sources of microbes and the interplay of dispersal with selection during initial colonization, as well as determine to what extent these findings are generalizable to primary and secondary succession in other ecosystems (e.g. volcanic soils, meteorites, post-fire recovery). These findings are timely considering the current unprecedented rates of glacier decline worldwide and the expansion of forelands. The microbial dynamics during the earliest stages of colonization and succession are critical to the eventual fate of these environments, as they develop into dynamic ecosystems harbouring biodiversity and driving globally-relevant biogeochemical cycles.

## Material and Methods

### Field sampling

Soil samples were collected from the forelands of two retreating glaciers: Sally Rocks tongue of Hurd Glacier on Livingston Island in Antarctica (August 2022) and Griessfirn Glacier in Canton Uri, Switzerland (March 2022). The Hurd Glacier is located at 62.6996°S, 60.4156°W, while the Griessfirn Glacier is at 46.84299°N, 8.82746°E. Specific coordinates for each sampling site at both glaciers are provided in **Table S1**. Previous studies at both sites were used to inform the sampling design including information on how deglaciation age was determined^28,52,80^. For the Antarctic site, nine surface soil samples (0-10 cm) were collected, corresponding to three soil age classes determined based on glacier front position measurements^81,82^: A (< 5 yrs), B (10 yrs), and C (21 yrs). Samples for 16S rRNA gene and metagenomic analyses were collected with a sterile spade, placed in whirl-pak sampling bags, frozen, and shipped to Monash University for processing. For Swiss samples, sampling locations were distributed along a well-defined soil chronosequence with increasing distance from the glacier terminus and categorized into four soil age classes: A (0-7yr), B (8-27yr), C (28-57yr) and D (58-127yr). In addition, at one location in each of the four soil-age classes, a 50-cm soil-profile was excavated to allow depth-resolved soil sampling at 10 cm intervals. Sampling locations were positioned along SW-NE-oriented transects that followed a single geomorphological landform, a band of lateral debris deposits, to minimize variability due to allogenic factors and intrinsic differences in microclimatic conditions and physical parameters among landform types^70^. Sampling was performed under dry-weather conditions and distant from previous rainfall. All samples were kept on ice or refrigerated until arrival in the laboratory and were thereafter stored at -20 °C (for molecular analysis) or 4 °C (for *ex situ* incubations) until further processing.

### Soil physicochemical analysis

Surface soils (0-10cm) from of the Swiss and Antarctic sites were analyzed at the Environmental Analysis Laboratory (EAL), Southern Cross University. In total, 37 separate soil physicochemical parameters were selected for analysis, based on commonly reported drivers of soil microbial composition globally. These included: phosphorus (mg/kg P), nitrate nitrogen (mg/kg N), ammonium nitrogen (mg/kg N), sulfur (mg/kg S), pH, electrical conductivity (dS/m), estimated organic matter (% OM), exchangeable calcium (mg/kg), exchangeable magnesium (mg/kg), exchangeable potassium (mg/kg), exchangeable sodium (mg/kg), exchangeable aluminium (mg/kg), exchangeable hydrogen (mg/kg), effective cation exchange capacity (ECEC) (cmol+/kg), calcium (%), magnesium (%), potassium (%), sodium - ESP (%), aluminium (%), hydrogen (%), calcium/magnesium ratio, zinc (mg/kg), manganese (mg/kg), iron (mg/kg), copper (mg/kg), boron (mg/kg), silicon (mg/kg Si), total carbon (%), total nitrogen (%), carbon/nitrogen ratio, chloride estimate (equiv. mg/kg), total organic carbon (%), moisture content (%), gravel (%), sand (%), silt (%), and clay (%).

### *In situ* gas and flux measurements

For the Swiss glacial chronosequence, *in situ* depth-resolved soil-gas samples were collected at 11 locations using a poly-use multilevel sampling system^71^. Details of its installation, sampling, and gas measurement procedures were described previously^83^. Concurrent with soil-gas sampling, we measured the depth-resolved volumetric soil-water content (cubic meters per cubic meter of soil) (PR2/6 capacitance probe; Delta-T Devices Ltd., Cambridge, UK) and recorded depth-resolved soil temperature (iButton temperature loggers; Maxim Integrated, San Jose, CA, USA) in 30-minute intervals throughout the duration of the sampling campaign. At 17 locations, we used static flux-chambers to quantify trace gas fluxes. Details of the installed chambers, sampling and measurement procedures were described previously^19,70^. In this study, the deployment of the collars at the new locations was performed prior to the start of the sampling campaign and included rainfall events to allow soil consolidation around the collars. Each flux chamber measurement lasted 90 min and consisted of nine gas samples collected at the following time intervals: four samples were collected within the first 10 min at nearly regular intervals, followed by five samples collected at increasingly longer intervals (10, 15, and 30 min). All gas samples were kept cooled until the measurement of trace gas concentrations by gas chromatography using a pulsed discharge helium-ionization detector (model TGA-6791-W-4U-2, Valco Instruments Company Inc.) as previously described^19^. Once the measurements were completed, topsoil samples were collected with a sterile spade at the flux-chamber locations for DNA extraction, quantitative polymerase chain reaction (qPCR) and metagenomic sequencing, and for *ex situ* incubations to quantify trace-gas oxidation rates using a previously described sampling procedure^19,71^. Finally, soil bulk-density measurements were performed at eight locations applying an adapted PU-foam method suitable for soils with high skeletal fraction^84^ as previously described^52^.

### *Ex situ* oxidation rates

To assess the capacity of the microbial communities at each glacier to oxidise trace gases (H_2_, CO, CH_4_), we placed the surface soils from the Swiss and Antarctic chronosequences and the soils from the Swiss depth profile samples in 120 ml serum vials. The wet weight of soil in each microcosm was recorded and used to normalise subsequent calculations. The headspace was continuously purged with air from a pressurized cylinder (Air Liquide) to achieve mixing ratios that reflect atmospheric levels, specifically 0.5 ppm H_2_, 0.6 ppm CO, and 1.8 ppm CH_4_. Sampling commenced immediately after sealing the vial and headspace samples of 2 ml were taken at five intervals. Heat-killed soils (subject to two autoclave cycles at 121 °C for 30 minutes each) and blank measurements using empty serum vials served as controls. These controls confirmed that trace gas oxidation in the samples was attributable to biotic processes. Gas concentrations were measured using the aforementioned gas chromatograph as previously described^19^. The power per cell derived from the oxidation of each trace gas was calculated as previously described^19,50^, by factoring in the reaction rate for each gas (based on *ex situ* oxidation rates at atmospheric concentrations), Gibbs energy of the reaction (based on standard Gibbs energy, the reaction quotient, gas-phase compound activities, and soil conditions), and the number of microbial cells involved (based on 16S rRNA gene copies quantified by qPCR and proportion of trace gas oxidising cells determined by metagenomics).

### Ammonium and sulfide oxidation

Oxic slurry experiments were undertaken to determine the oxidation rates of ammonium and sulfide in surface soils collected from both glaciers. Sterilised surface soils (autoclaved at 120°C for 1 hour) were used as controls to confirm that the observed oxidation rates were driven by solely biotic processes. Slurries containing 10 g soil (wet weight) and 200 mL substrate amended-ultrapure water (50 µM of either ammonium chloride or sodium sulfide) were prepared in 250 mL Schott bottles. The slurries were aerated for 5 minutes to ensure oxic conditions. The bottles containing the slurries were left uncapped but loosely covered with pre-combusted aluminium foil. The slurries were incubated in the dark and were mixed from time to time for the duration of the incubation period (up to 22 days). At each time point, 15mL of samples were collected and filtered through 0.22 µm pore-sized filters (Sartorius Minisart syringe filter). Filtered samples were analyzed for ammonium, nitrate, nitrite and sulfate concentrations using a Lachat Quickchem 8000 Flow Injection Analyzer following the procedures in Standard Methods for Water and Wastewater (APHA 2005)^85^. Oxidation rates of ammonium were calculated using linear and non-linear regression of ammonium consumption over time, as well as linear regression of nitrite and nitrate accumulation over time. Similarly, rates of sulfide oxidation were calculated from linear regression of sulfate increase over time.

### Chlorophyll *a* measurement

To extract chlorophyll from each sample, 9 ml of acetone was added to 5 g of soil, mixed, sonicated in ultrasonic water bath at a maximum setting for 5 minutes, and incubated overnight in refrigerator. Then, 1 ml of RO water was added to each sample and these were subsequently centrifuged for 10 minutes at 2,000 rpm. 3 ml of liquid phase of sample extract were transferred to a cuvette and chlorophyll was measured on a spectrophotometer (Eppendorf BioSpectrometer) with wavelength scan at 665 and 750 nm. Lastly chlorophyll *a* concentration was measured with the formula:

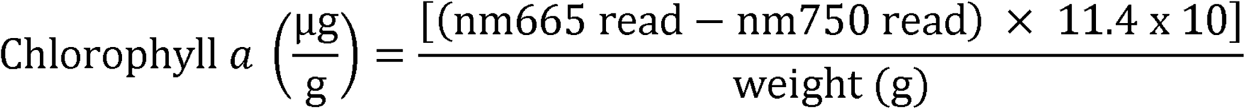

### Community DNA extraction

Total community DNA was extracted for each of the Swiss and Antarctic samples using 0.5 g of soil. Extractions were performed using the MoBio PowerSoil Isolation kit according to the manufacturer’s instructions, with samples eluted in DNase- and RNase-free UltraPure Water (ThermoFisher). A sample-free negative control was also run for each glacier sampled. Nucleic acid purity, yield, and integrity were measured using a NanoDrop ND-1000 spectrophotometer, a Qubit Fluorometer 2.0, and through agarose gel electrophoresis.

### Quantitative PCR

Quantitative polymerase chain reactions (qPCR) were used to estimate total bacterial and archaeal biomass of biocrust and topsoil samples. The 16S rRNA gene was amplified using the degenerate primer pair (515 F 5 -154 GTGY-CAGCMGCCGCGGTAA-3) and 806R 5 -GGACTACNVGGGTWTCTAAT-3). A synthetic *E. coli* 16S rRNA gene sequence in a pUC-like cloning vector (pMA plasmid; GeneArt, ThermoFisher Scientific) was used as a standard. PCR reactions were set up in each well of a 96-well plate using LightCycler 480 SYBR Green I Master Mix. Each sample was run in triplicate and standards in duplicate on a LightCycler 480 Instrument II (Roche). The qPCR conditions were as follows: pre-incubation at 95 °C for 3 min and 45 cycles of denaturation 95 °C for 30 s, annealing at 54 °C for 30 s, and extension at 72 °C for 24 s. 16S rRNA gene copy numbers were calculated based on a standard curve constructed by plotting average C_p_ values of a serial dilution of the plasmid-borne standard against their copy numbers.

### Community profiling

16S rRNA gene amplicon sequencing was used to infer the composition of the bacterial and archaeal community in each sample. Specifically, the V4 hypervariable region of the 16S rRNA gene was amplified using the universal primer pairs F515 and R806 from the Earth Microbiome Project^86^. Illumina paired-end sequencing (2 × 300 bp) was conducted at the Australian Centre for Ecogenomics, University of Queensland. The resulting raw sequences underwent quality filtering, primer trimming, denoising, and removal of singletons within the QIIME 2 platform^87^. The final dataset comprised 585,111 high-quality 16S rRNA amplicon sequence variants (ASVs).

### Biodiversity analyses

Alpha and beta diversity were calculated using R v 4.1.0 and the Phyloseq package^88^. First, reads were normalized using the coverage-based rarefaction and extrapolation method implemented in the R package iNEXT^89^. Coverage was calculated for each sample using the function phyloseq_coverage, followed by rarefaction of all reads, using the function phyloseq_coverage_raref with default parameters. Observed and estimated richness was calculated using the estimate_richness function specifying “observed” and “chao1” flags. To calculate beta diversity, sample counts were first transformed to relative abundance and clustered using a Non-metric Multidimensional Scaling ordination via the ordinate with flags “NMDS” and “Bray” function. A stress plot was used to determine linear (R^2^=0.839) and non-metric (R^2^=0.967) fit between ordination distance and dissimilarity (Bray-Curtis). The final stress of <0.2 (0.182) indicated a good representation in reduced dimension. To test for significant difference in microbial community structure between soil age and location a Permutational Multivariate Analysis of Variance Using Distance Matrices was used via the adonis function using the R package Vegan^90^. Finally, to determine if significant results were because of dispersion a Multivariate homogeneity of groups dispersions test was used via the betadisper function. The multisite metric zeta diversity (ζ) was used to assess incidence-based turnover in community composition (ASVs) along the Swiss glacier foreland using the *zetadiv* R package^63^. ζ_2_ and ζ_4_ distance decay were computed via the *zeta.ddecay* function on a Jaccard-normalized presence-absence ASVs dataset with subsampling set to 1000. Zeta decline was calculated with the function *zeta.decline.mc*, performing subsampling set to 1000 on a Jaccard-normalized presence-absence ASVs dataset for ζ orders ζ_1_ to ζ_6_. This approach captured community structuring across the foreland, with increasing ζ values approaching zero. Power-law and exponential models were fitted to the ζ decline curves, using Akaike Information Criterion (AIC) scores to evaluate which model best explained the relationship between ζ diversity and order *i*. Zeta variation partitioning was performed via the function *zeta.varpart* on a presence-absence ASVs dataset using *method.glm* = *glm.fit2*. To determine the optimal model for *zeta.varpart*, we used redundancy analysis computed via the *rda* function in the *vegan* R package (REF), retaining only non-collinear physicochemical variables.

### Metagenomic sequencing and metabolic profiling

For the Swiss samples, metagenomic shotgun libraries were prepared using the Nextera XT DNA Sample Preparation Kit (Illumina Inc., San Diego, CA, USA) and subject to paired-end sequencing (2 × 150 bp) on an Illumina NovaSeq6000 platform at the Australian Centre for Ecogenomics (ACE), University of Queensland. For the Antarctic samples, metagenomic shotgun libraries were prepared and subject to paired-end (2 × 100 bp) on a DNBSEQ-G400 platform at Micromon Genomics, Monash University.

### Metagenomic short read analysis

To estimate the metabolic capability of the soil communities, quality-filtered short reads were searched against custom protein databases of 51 metabolic marker genes (https://bridges.monash.edu/collections/Greening_lab_metabolic_marker_gene_data bases/5230745) covering major pathways of carbon fixation, phototrophy, aerobic and anaerobic respiration, fermentation, and nitrogen, sulfur, iron, hydrogen, carbon monoxide, and methane cycling^91–93^. DIAMOND v.0.9.31 searches^94^ were conducted with a query coverage of 80% and identity threshold of 50%, except for *rho* (40%), *nuoF,* group 4 [NiFe]-hydrogenases*, mmoX,* [FeFe]-hydrogenases*, coxL, amoA, nxrA, rbcL* (all 60%), *psaA* (80%), *psbA, isoA, atpA, aro, ygfK* (70%), *and hbsT* (75%). Read counts for each gene were normalized to reads per kilobase per million (RPKM) and average gene copy per organism as previously described^19,50,95^.

### Metabolic gene driver analyses

Generalized linear models (GLMs) were used to determine the effects of soil age and depth on metabolic gene abundance based on metagenomic short reads. Following mean – variance comparisons to detect overdispersion and comparing optimal model fits against the residual plots of each distribution, GLMs were fitted with a Gaussian distribution. Other important environmental predictors were assessed using a random Forest analysis via the R package randomForest^96^. Collinearity between predictor variables was assessed by constructing Pearsons’s correlation matrices using the *cor* function in base R, between the 37 soil physicochemical components measured in this study. Variable reduction was conducted through an ecologically informed iterative process. Initially, correlations among potential predictors were calculated. Highly correlated pairs (r ≥ 0.9) were manually inspected. Following each correlation analysis, the less ecologically significant variable of a highly correlated pair was removed, based on established knowledge of key environmental drivers of microbial communities in aerated soil ecosystems^97^. For instance, if zinc and pH were found to be highly correlated, pH would be retained due to its recognized overriding ecological importance. This process was repeated to refine the set of variables, resulting in a final set of predictors which were used for subsequent analyses. A random forest model was then generated assessing the importance of each predictor against trace gas oxidation rates, using the *randomForest* function. Number of variables randomly sampled as candidates at each split = 10, number of trees grown = 10,000 and sampling of cases was conducted with replacement.

### Metagenome-assembled genome analysis

Using the Metaphor pipeline^98^, the metagenomes of the Antarctic and Swiss datasets were co-assembled independently of each other using MEGAHIT v1.2.9^99^ with default settings. Contigs shorter than 1,000 bp were discarded. The assembled contigs were binned with Vamb v4.1.3^100^, MetaBAT v2.12.1^101^, CONCOCT v1.1.0^102^ and SemiBin2^103^. The four bin sets were subsequently refined with MetaWRAP’s refinement module^104^ and de-replicated with dRep v3.4.2^105^ with 95% ANI integrated with CheckM2^106^. The completeness and contamination of the bins were assessed using CheckM2^106^. Quality thresholds for bins were selected based on a previous study^107^, retaining only medium (completeness >50%, contamination <10%) and high (completeness >90%, contamination <5%) quality bins for further processing, which were termed metagenome-assembled genomes (MAGs). MAGs taxonomy was determined using the Genome Taxonomy Database Release R214^108^ via GTDB-Tk v2.3.2^109^. Proteins from each MAG were predicted intrinsically in CheckM2^106^. The MAGs were metabolically annotated against the above-described custom database using DIAMOND v.0.9.31. Gene hits were filtered to retain only those either at least 40 amino acids in length or with at least 80% query or 80% subject coverage. For predicted proteins, the same identity thresholds were used as above except for *atpA* (60%), *psbA* (60%), *rdhA* (45%), *cyc2* (35%), and *rho* (30%).

### Phylogenetic analysis

The genome phylogenetic tree, comprising high- and medium-quality MAGs (completion >50% and contamination <10%) bacterial and archaeal MAGs, was constructed using PhyloPhLan v3.0.67^110^. Multiple sequence alignment was performed using MAFFT v7.508^111^ by identifying 400 universal proteins from these microbial genomes, following Segata et al. (2013)^112^.The tree was inferred with IQ-TREE v. 2.2.0.3^113^ using model LG+F+G4 for and 1,000 ultrafast bootstrap iterations. The tree was midpoint rooted and rendered in iTOL v7^114^.

### Habitat specialization indices

The degree of habitat specialization of each MAG (i.e. whether they were relative ‘habitat generalists’ or ‘habitat specialists’) was calculated based on the coefficient of variance in the metagenomes (based on read mapping). The mean and standard deviation community specialization index was 1.459 ± 0.621 for the Antarctic MAGs and 1.302 ± 0.792 for the Swiss MAGs. Relative ‘habitat generalists’ were defined as MAGs with a specialization index (i.e. coefficient of variance) below the first quartile (below 0.9886 for Antarctic MAGs, 0.6652 for Swiss MAGs) and relative ‘habitat specialists’ were defined as MAGs with a specialization index in the fourth quartile (above 1.8751 for Antarctic MAGs, 1.7778 for Swiss MAGs). To disentangle the effects of soil age and depth for the Swiss samples, habitat specialization index was only calculated based on read mapping to the surface metagenomes. Using the 16S rRNA gene amplicon sequencing data, the habitat specialization indices of each genus and phylum with an average relative abundance exceeding 0.005% was also calculated, based on their coefficient of variance (habitat generalists and specialists defined as taxa with specialization indices below the first quartile and above the fourth quartile respectively).

## Acknowledgements

This study was supported by an NHMRC EL2 Fellowship (APP1178715; to C.G.), the Human Frontiers Science Program (RGY0058/2022; to C.G. and J.A.B.), ARC Discovery Early Career Research Award (DE230101346; to S.K.B., DE250101210; to P.M.L.), a Monash University Early Career Postdoctoral Fellowship (to F.R.), an Australian Research Council SRIEAS Grant SR200100005 Securing Antarctica’s Environmental Future (to C.G.), a Swiss National Science Foundation Early Mobility Postdoctoral Fellowship (to E.C.), NERC (NE/T010967/1), the Agence Nationale de la Recherche (ANR23-CPJ1-0172-01) and the European Research Council (ERC) under the European Union’s Horizon Europe Research and Innovation programme (Grant agreement No. 101115755) (to J.A.B), and a Agencia Estatal de Investigación grant (PID2019-105469RB-C22; to A.d.l.R.). We thank Steven L. Chown for helpful discussions.

## Author contributions

E.C., C.G., P.A.N., and M.H.S. conceived and designed this study. C.G., M.H.S., and P.L.M.C. supervised this study. E.C., P.A.N., M.H.S., B.F.M. and A.dl.R. conducted fieldwork. E.C., S.K.B., P.A.N., W.W.W., L.J., T.J., G.N., and M.H. conducted and analysed biogeochemical assays. S.K.B., F.R., C.G., and E.C. conducted community and biodiversity analyses. F.R., C.G., G.N., S.K.B., and P.M.L. conducted metagenomic analyses. J.A.B. and A.S. provided ecological insights. F.R. and C.G. wrote the paper with input from all authors.

## Data availability

All sequence data from this study is available at the Sequence Read Archive, with accession numbers PRJNA1178459 for metagenomic sequences, PRJNA1178814 for 16S rRNA gene sequences, PRJNA1206715 for Antarctic MAGs, and PRJNA1206714 for Swiss MAGs.

## Supplementary Figures

**Supp. Figure 1.**
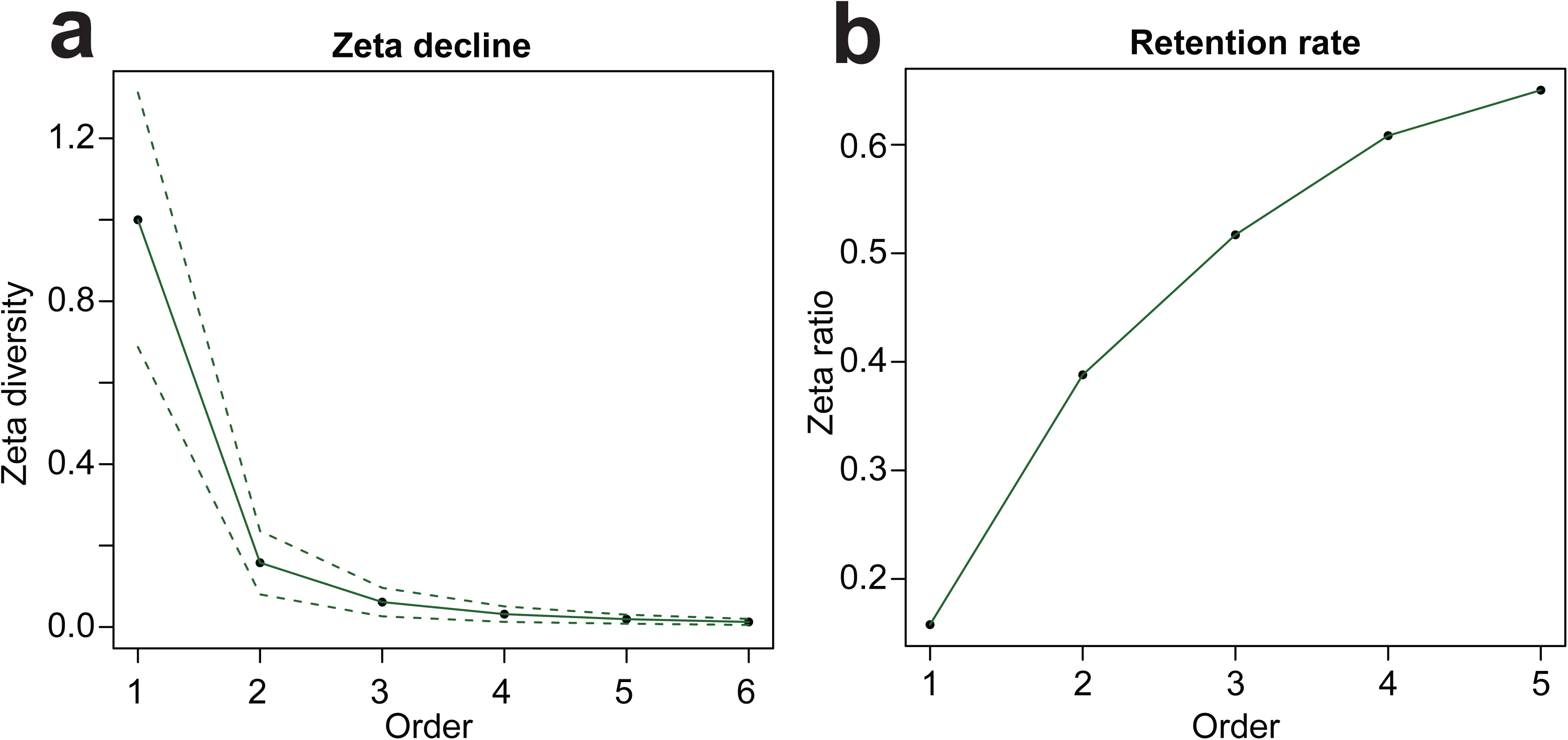
Multisite community turnover and assembly processes along the Swiss glacial foreland. **(a)** Zeta decline illustrates how the number of shared taxa (ASVs) decreases as more sites are included in the comparison (Zeta Order). **(b)** Zeta diversity ratio showing the rate of taxon retention, highlighting the probability of retaining common taxa over rare ones at any given order as additional sites are included in the comparison.

**Supp. Figure 2.**
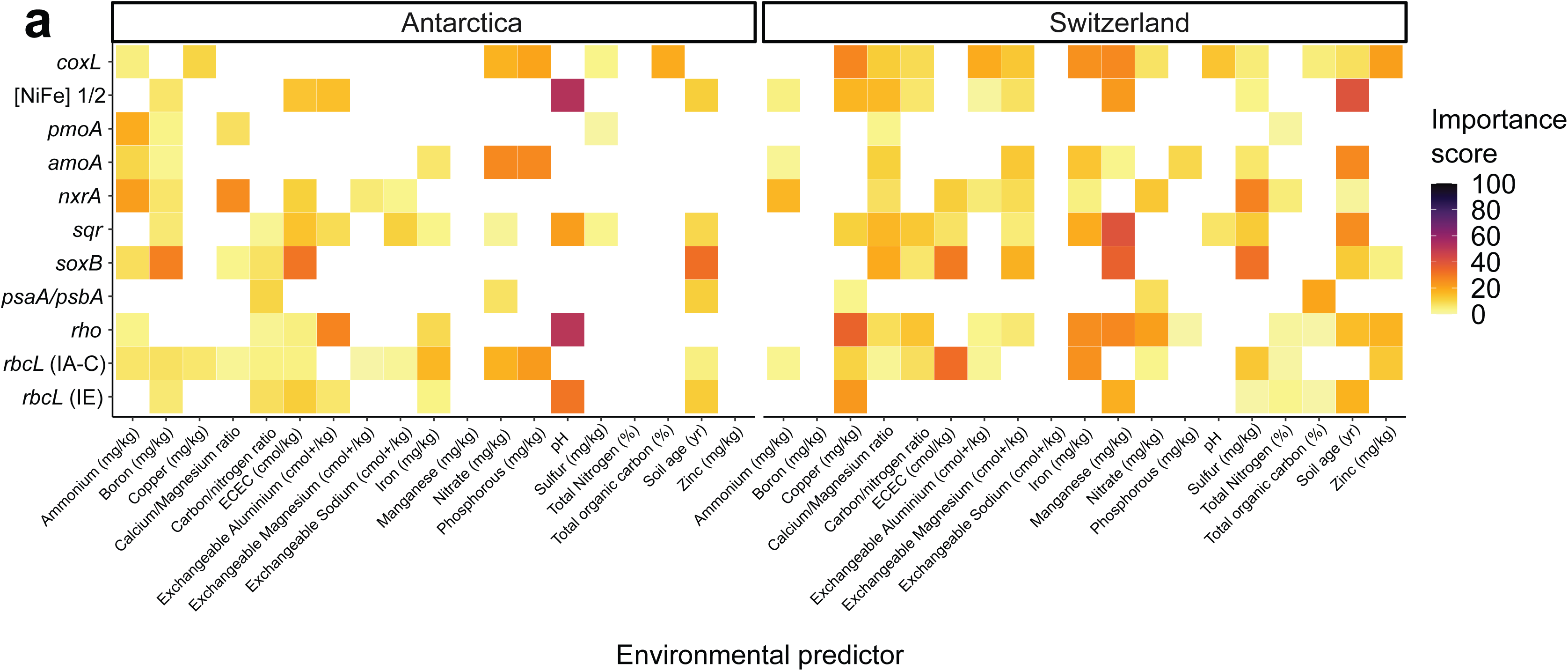
Random forest analysis showing the influence (importance score) of environmental variables over the relative abundance of metabolic marker genes for trace gas oxidation, nitrification, sulfide and thiosulfide oxidation, phototrophy and carbon fixation recovered from the Antarctic and Swiss glacial forelands.

**Supp. Figure 3.**
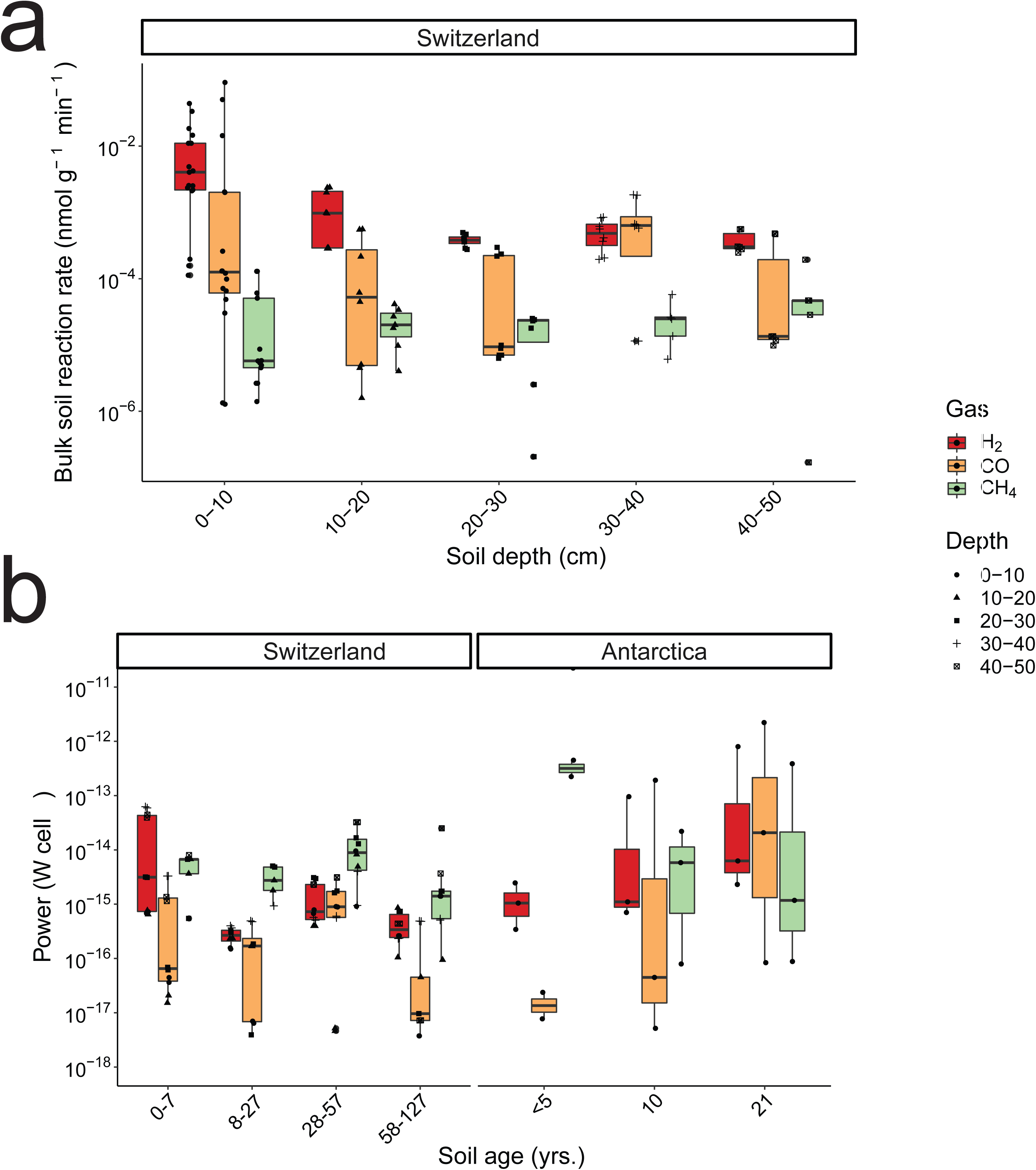
**(a)** Bulk soil oxidation rate for each gas, according to soil depth of Swiss samples with shapes showing respective soil age. Measurements were performed with at least four biological replicates per age group. **(b)** Amount of power per cell generated from the oxidation of each trace gas, calculated using thermodynamic models based on ex situ rates measured on five depths for the Swiss glacial foreland and on top Antarctic foreland soil. Measurements were performed with at least two biological replicates per age group.

## Supplementary Tables

**Supp. Table 1.** a, Samples metadata. b, Soil physicochemical parameters measured along the Antarctic and Swiss glacial forelands.

**Supp. Table 2.** a, DNA concentration, elution volume, sample amount, and 16S copy number normalized to both wet weight and dry weight are shown for each sample used in metagenomic sequencing, including extraction controls. b, Amplicon sequence variant (ASV) table showing quality trimmed counts for each sample. c, Relative abundance summary at phylum level. d, lpha diversity (Shannon) for each sample. e, Beta diversity, testing for significant differences in composition between location and soil age through PERMANOVA. f, Chlorophyll a measurement for each sample.

**Supp. Table 3.** a, Antarctic glacial foreland short read annotation. b, Swiss glacial foreland short read annotation. c, Functional gene abundance driver analysis using Generalised Linear Model. d, Functional gene abundance driver analysis using Random Forest analysis. e, Antarctica glacial foreland MAG annotation. e, Switzerland glacial foreland MAG annotation. g, specialization index cutoffs.

**Supp. Table 4.** a, Antarctic MAGs relative abundance and specialization index. b, Swiss MAGs relative abundance and specialization index. c, Antarctic and Swiss generalist, intermediate and specialist MAGs proportion. d, Antarctic metabolic modules. e, Antarctic gene count. f, Antarctic metabolic modules across generalist, intermediate and specialist MAGs. g, Swiss metabolic modules. h, Swiss gene count. i, Swiss metabolic modules across generalist, intermediate and specialist MAGs.

**Supp. Table 5.** a-d, Gas rates and fluxes. a, Rates and power calculations. b, Flux calculations. c, Nitrification and sulfide oxidation rates with linear and non-linear models fitted to each curve and best rates picked based on AIC. d, Rates and Power legend.

